# Magnitude and Correlates of Caesarean Section in Urban and Rural Areas: A Multivariate Study in Vietnam

**DOI:** 10.1101/554964

**Authors:** Myriam de Loenzien, Clémence Schantz, Bich Ngoc Luu, Alexandre Dumont

**Author notes:** These authors contributed equally to this work (Clémence Schantz, Bich Ngoc Luu, Alexandre Dumont).

## Abstract

Caesarean section can prevent maternal and neonatal mortality and morbidity. However, it involves risks and high costs which can be a burden, especially in low and middle income countries. The international healthcare community considers the optimal caesarean rate to be between 10% and 15%. The aim of this study is to assess its magnitude and correlates among women of reproductive age in urban and rural areas in Vietnam. We analyzed microdata from the national Multiple Indicator Cluster Survey (MICS) conducted in 2013-2014 using representative sample of households at the national level as well as regarding the urban and the rural areas. A total of 1,378 women who delivered in institutional settings in the two years preceding the survey were included. Frequency and percentage distributions of the variables were performed. Bivariate and multivariate logistic regression analysis were undertaken to identify the factors associated with caesarean section. Odds ratios with 95% confidence interval were used to ascertain the direction and strength of the associations. The overall CS rate among the women who delivered in healthcare facilities in Vietnam is particularly high (29.2%) with regards to WHO standards. After controlling for significant characteristics, living in urban areas more than doubles the likelihood of undergoing a CS (OR = 2.31; 95% CI 1.79 to 2.98). Maternal age at delivery over 35 is a major positive correlate of CS. Beyond this common phenomenon, distinct lines of socioeconomic and demographic cleavage operate in urban versus rural areas. The differences regarding correlates of CS according to the place of residence suggest that specific measures should be taken in each setting to allow women to access childbirth services appropriate to their needs. Further research is needed on this topic.

## Introduction

Caesarean section can prevent maternal and neonatal mortality and morbidity. However, it involves risks and high costs which can be a burden, especially in low and middle income countries. The international healthcare community considers the optimal caesarean rate to be between 10% and 15% (1). Urbanization, which is related not only to a population moving from a rural to an urban area and an increased concentration of people living in urban areas but also to the whole process of societal adaptation to subsequent changes, has been identified as a prominent contributing factor to caesarean section (CS) practices in several countries and areas (2)(3)(4)(5)(6)(7)(8). However, this influence is controversial (9)(10)(11).

Vietnam, which transformed from a low to a middle income country in the last decade, has witnessed increasing CS rates concomitantly with urbanization. In this country, the proportion of women undergoing CS increased from 3.4% in 1997 (12) to 27.5% in 2014 (13), which largely exceeded the levels recommended by the World Health Organization (WHO) (10 to 15%) (14). This percentage is among the highest in the region (14)(15), and this trend shows no sign of abatement. The increase is occurring in a context of rapid socioeconomic and demographic changes. During the same period, the proportion of people living in urban areas rapidly increased from 23.7% in 1999 to 29.6% in 2009 (16).

We propose to measure the influence of living in urban versus rural areas on childbirth practices and to explore the possible pathways of the influence of the place of residence on CS in Vietnam. Using microdata from a nationally representative sample, we provide the sociodemographic profile associated with high CS rates. Subsequently, we present correlates of CS rates by making a distinction between women who live in rural and urban areas. Our main argument is that beyond the apparent overall convergence, CS practices diverge not only in magnitude between rural and urban areas but also regarding their dynamic.

## Literature review

### Relationship between CS and place of residence

For several decades, living in urban areas in low- and middle-income countries in Asia, Africa and Latin America has been associated with higher CS rates after controlling for multiple socioeconomic, biomedical and institutional factors (2)(3)(4)(5)(6)(7)(8). However, this relationship appears to be nonsignificant in various settings (9)(10). Some studies have even shown a reverse trend. In Hawaii, despite a lower risk of delivery by CS, women who deliver in rural hospitals have higher rates of primary CS than do women who deliver in urban hospitals, even after controlling for maternal risk factors (11).

Further analyses taking into account the level of urbanization complement these results. In Taiwan, CS rates increased with an advancing urbanization level (17). Similarly, a study using data from 29 countries in Asia, Africa and Latin America showed higher CS rates in urban areas than in periurban areas (18). Conversely, a study in Cambodia showed that CS rates were lower for women living in Phnom Penh than for women living in its surrounding area (19).

Some studies go more in depth by conducting intersectional analyses between the place of residence and the wealth effect. One analysis that adopted data from demographic and health surveys performed in low- and middle-income countries in Africa, Asia and Latin America showed that the CS rates in most countries were higher in urban areas than in rural richer households, which represented half of the rural population. In turn, most rural richer households had higher CS rates than did rural poorer households (20).

More refined indicators of childbirth practices have also been used. In more-developed countries, research indicates a higher level of non-medically indicated labor induction in urban areas but a more rapid rise in rural areas, such as the trends observed in the United States (21). In states in Burkina Faso, CS deliveries for nonabsolute medical indications were more frequent among women living in urban areas even after controlling for other factors (22).

In this study, we consider that the decision to undergo caesarean section results from a negotiation between the caregiver and the patient, which is determined by proximate determinants. Among them, patient’s and health caregivers’ perceptions play major roles, as well as the availability and accessibility of healthcare facilities, equipment, personnel and technologies (23). These proximate determinants are in turn determined by distal determinants, such as biomedical factors, but also social, cultural and political characteristics at individual, interindividual and collective levels. These characteristics include women’s human, economic and social capital but also cultural beliefs, values and norms regarding family and gender relations (24), interactions between social groups (9), the media and formal institutions, welfare state and national policies as well as economic conditions (3)(25). Due to contrasted modes of socialization and levels of equipment, we expect underlying processes related to these phenomena to differ between rural and urban areas.

### CS and urbanization in Vietnam

In Vietnam, urbanization has rapidly developed and continues to exhibit a rising trend in the context of economic growth and on-going demographic transition. Urbanization accelerated in the 1990s (26) following the reforms in the mid-1980s from a centralized system to a market-oriented economy under state guidance (27). The country has shifted its policy from the promotion of intermediate-level cities in the 1990s-2000s to more investment in great metropolitan areas aimed at acting as drivers of the economy (28). The urbanization rate increased from 19.2% in the 1980s to 29.6% in 2009 and reached 34.0% in 2015 (16)(29). Simultaneously, rural-urban inequalities have decreased (30). This reduction has mainly been due to migration in a context of improving economic conditions, the development of industrialization, increasing international integration and profound demographic and technological changes (31).

Overall, the proportion of women who deliver by CS has multiplied by almost 7 within a 17-year period. From a very low level in 1997 (3.4%), this proportion reached the level proven to be the threshold of the absence of efficiency of CS (10%) in 2002 (9.9%) (12) (32) (14). This rate reached almost three times this value in 2013-14 (27.5%) (33)(13).

CS rates have increased at a higher pace in rural areas, where they have multiplied by 9 (from 2.3% to 21.0%), than they have in urban areas, where they have multiplied by 3 (from 13.6% to 43.3%). Consequently, the urban-rural ratio of the proportion of women who underwent CS dropped (from 5.9 in 1997 to 2.1 in 2014). This increase in CS rates occurred in the context of a marked development of childbirth biomedicalization fostered by recent investment in district hospitals by the government (34).

Whereas only a minority of pregnancies were followed up by a doctor in 1997 (28.2%), almost all pregnancies were followed up in 2014 (90.3%). The number of prenatal care visits has dramatically increased. Very few women attended 7 visits or more in 1997 (2.3%), and a higher proportion of women utilized this number of visits 17 years later (39.0%). As a result, neonatal mortality rates dropped (from 20. to 11.4 deaths per 1000 live births), as did maternal mortality rates (from 100 to 54 maternal deaths per 100,000 live births) (34). Vietnam ended its demographic transition, with fertility reaching the replacement level since 2005 (35).

Studies in this country suggest that urban areas are linked to a higher level of CSs. However, these studies referred to a period when CS rates were still low (9) or to specific geographical areas (36). Therefore, there is a need to update general trends in this country and to better understand the correlates of such difference between rural and urban areas.

## Materials and Methods

### Data

We used data from the 2013-2014 Multiple Indicator Clusters Survey (MICS). Urban and rural areas within each region were identified as the main sampling strata. The sample was selected in two stages: census enumeration areas were selected within each stratum, and households were selected within each enumeration area (13). This dataset provides statistically representative samples of women aged 15-49 years at the national and regional levels and for each type of setting (rural versus urban). Among these datasets, we focus on women who had a singleton birth at least once during the 2 years prior to the survey. This population represents 1,453 women. Among these women, we take into account those who delivered at an institutional setting, accounting for 94.4% of the population. Only the last birth of each woman was considered.

### Outcome measures and covariates

The outcome variable for this study is the woman’s mode of delivery, either through CS or vaginal delivery.

The main covariate for this study is the place of residence (urban versus rural area). The influence of this variable is explored by controlling for other socioeconomic and demographic correlates. We take into account the place of delivery (public versus private health sector). Based on empirical observations of the distribution, the number of antenatal care visits was distinguished between 6 visits or fewer versus 7 or more to maximize the rural-urban difference. We also took into account the birth weight of the newborn as perceived by the mother (less than 2.8 kg, 2.8 to 3.5 kg, 3.6 kg and over), the maternal age at delivery (15 to 19 years, 20 to 34 years or 35 years and over) and the woman’s past experience of childbirth (primiparous versus multiparous). Due to the preference for sons associated with CS in Vietnam, we also explored the influence of the sex of the newborn (37). Additional sociodemographic and cultural characteristics included the level of education of the women (primary or less, secondary or tertiary), the region (North Central, Mekong River delta, Red River delta, Northern Midlands, Central Highlands or the Southeast), the quintile of wealth of the household (poorest, poor, middle, rich or richest), and the ethnicity of the household head (Kinh ethnic group versus minority ethnic groups).

### Analysis

We conducted bivariate analysis and stepwise logistic regression to assess the characteristics associated with CS practice as opposed to vaginal delivery. Multivariate logistic regression models allowed for comparisons between models for all women and for only women living in rural or urban areas. For each of these 3 groups, two models were tested. The first model (restricted model) included only sociodemographic variables, whereas the complete model took into account all the available variables, including sociocultural characteristics. Bivariate analyses used the women’s sample weights. The multivariate results included all variables that reached the minimum level of significance in the bivariate analysis. These results are controlled for the cluster effect. For each model and for the chi-square tests, we draw on two levels of risk (p < 0.05 and p < 0.10). All statistical analyses were performed with IBM PASW Statistics 18 software at Paris Descartes University.

### Compliance with ethical standards

The Vietnam General Statistics Office (GSO) and the United Nations Children’s Fund (UNICEF) approved the tools of the Vietnam Multiple Indicator Cluster Survey (MICS) before the survey was conducted, in accordance with the ethical standards laid down in the 1964 Declaration of Helsinki and its later amendments or comparable ethical standards. Participation was voluntary, and informed consent was obtained from all the individual participants included in the study. The MICS data are freely available through the UNICEF MICS website, and there is no need to obtain ethical approval before using the data. To access data from the MICS website, a written request was submitted to UNICEF, and permission was granted.

## Results

Table 1 provides an overview of the social and demographic profiles of the women who delivered in healthcare facilities and the corresponding rates of CS.

**Table 1.**
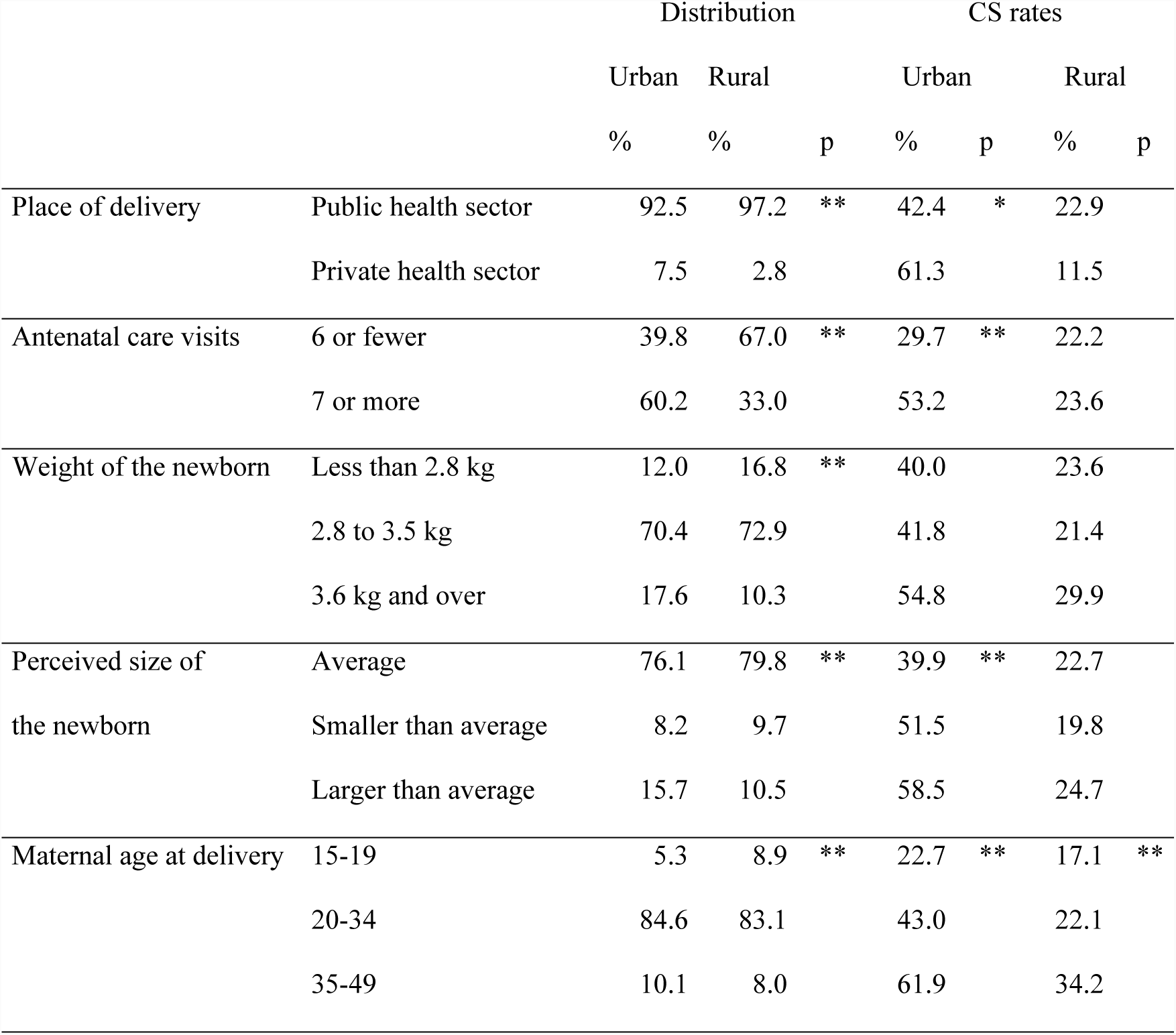
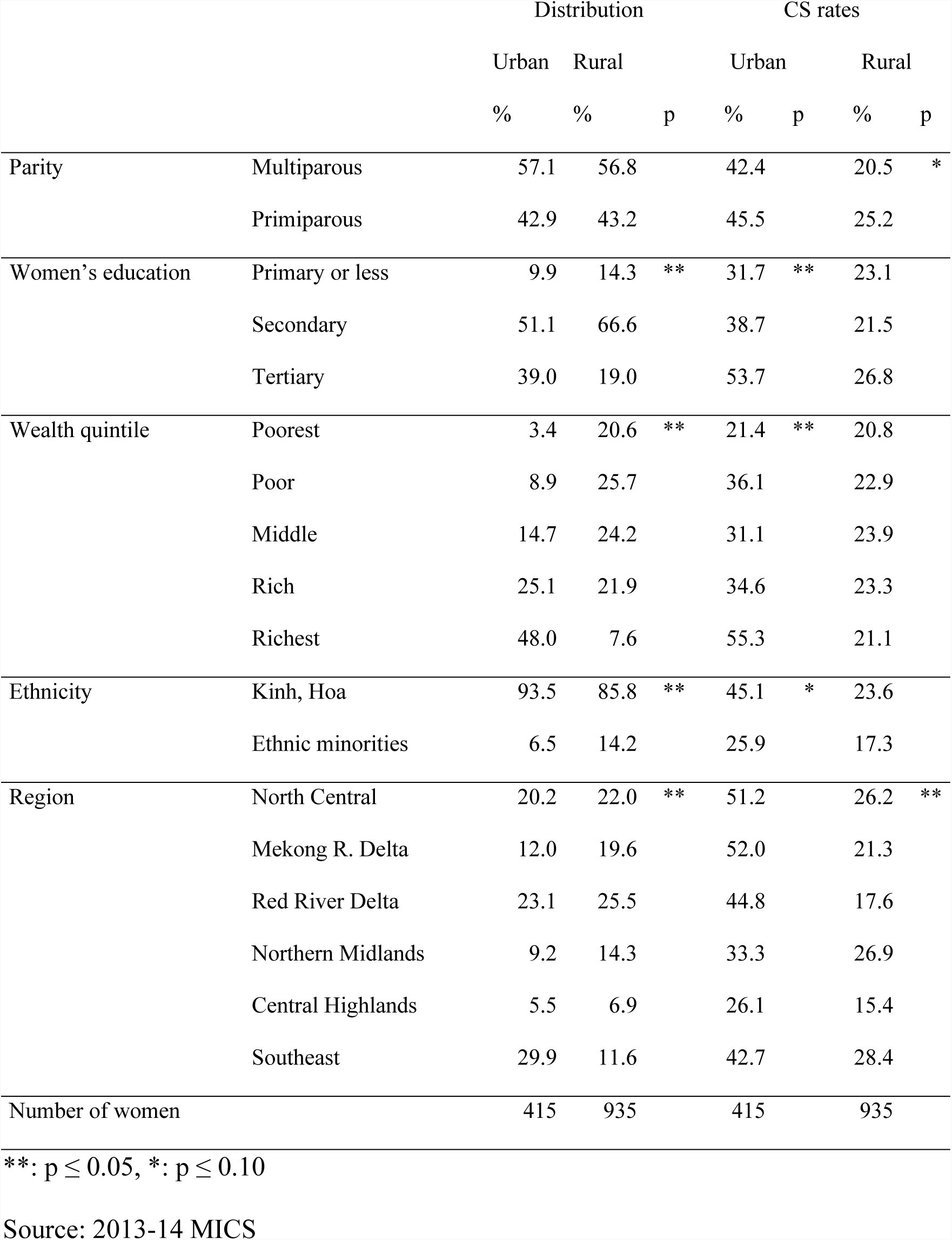
Social and demographic profiles of the women who had a singleton birth in the two years preceding the survey in healthcare facilities and corresponding rates of caesarean section (CS) (n = 1350 deliveries)

Overall, almost one-third of the women live in an urban area (30.7%). Several correlates linked to higher levels of CS are more prevalent in the urban areas than in the rural areas. First, among the women who deliver in institutional settings, almost two-thirds of those living in urban areas have more than 7 antenatal care visits, whereas this is the case for only one-third of those living in rural areas. Second, several indicators show more favorable socioeconomic situations for women living in urban areas: a much larger proportion of women reach a tertiary level of education in urban (39.0%) than in rural areas (19.0%), the proportion of women in the richest household quintile reaches a much higher level in urban areas (48.0%) than in rural areas (7.6%), and lower proportions of women belonging to minority ethnic groups are observed in urban (6.5%) than in rural areas (14.2%). Third, the maternal age at delivery is lower in rural areas than in urban areas. The proportion of women who deliver after 35 is higher in urban (10.1%) than in rural areas (8.0%) and conversely, fewer women deliver between 15 and 19 in urban (5.3%) than in rural areas (8.9%).

The overall CS rate among the women who delivered in healthcare facilities is particularly high (29.2%) with regards to WHO standards (14). The CS rate is almost twice as high in urban (42.4%) than in rural areas (22.9%). The results regarding CS rates confirm that the urban context is particularly favorable to CS. First, in urban areas, CS rates were almost doubled among women who had at least 7 antenatal care visits compared to those among women with 6 visits or fewer. Second, in urban areas, women who have a higher level of education have higher CS rates, those who live in the richest households also have higher CS rates, and to a lesser extent those who belong to the Kinh ethnic group have higher CS rates than those belonging to the minority ethnic groups. Third, a higher maternal age at delivery is associated with higher CS rates in both rural and urban areas.

A higher number of antenatal care visits, higher levels of education, wealth, and concentration of Kinh ethnic groups and higher maternal age at delivery in urban areas combined with higher CS rates among people of these groups help to understand part of the urban–rural gap regarding CS rates. However, more in-depth analysis is needed to document the relative influence of each of these correlates on the urban–rural difference in CS rates, which will be achieved using multivariate analysis.

The results of the analysis of correlates of CS for the whole population are displayed in Table 2. We will first examine a model taking into account only demographic and medical variables. We will subsequently examine a model that also includes the socioeconomic variables. The results show that after controlling for significant characteristics, living in urban areas more than doubles the likelihood of undergoing a CS (OR = 2.31; 95% CI 1.79 to 2.98, see restricted model). The completed model shows that the influence of the place of residence on CS weakens when ethnicity is taken into account. This is partly due to the higher concentration of population belonging to minority ethnic groups in rural areas. Following the same trend, the weakening of the positive influence of having 7 antenatal care visits or more suggests that the higher level of medicalization in urban areas mostly regards women from the Kinh ethnic group. In addition to the place of residence, the number of antenatal care visits and ethnicity, delivering at 35 years or over remains strongly linked to CS, as is also the case of maternal perception of the newborn weight as above average and delivery for the first time.

**Table 2.** Multivariate analysis of the factors associated with caesarean delivery (n = 1350)^a^.

|  | Restricted model |  |  | Complete model |  |  |
| --- | --- | --- | --- | --- | --- | --- |
|  | OR | 95% CI |  | OR | 95% CI |  |
| Place of residence (reference = rural) |  |  |  |  |  |  |
| Urban | 2.31** | 1.79 | 2.98 | 1.99** | 1.48 | 2.67 |
| Place of delivery (reference = public sector) |  |  |  |  |  |  |
| Private sector | 1.35 | 0.82 | 2.21 | 1.27 | 0.76 | 2.13 |
| Antenatal care visits (reference = 6 or fewer) |  |  |  |  |  |  |
| 7 visits or more | 1.38** | 1.06 | 1.79 | 1.32 | 0.98 | 1.77 |
| Weight of newborn (reference = 2.8 to 3.5 kg) |  |  |  |  |  |  |
| Less than 2.8 kg | 1.05 | 0.72 | 1.52 | 1.08 | 0.74 | 1.59 |
| 3.6 kg and over | 1.60** | 1.10 | 2.32 | 1.55** | 1.06 | 2.26 |
| Maternal age at delivery (reference = 20-34) |  |  |  |  |  |  |
| 15-19 | 0.55 | 0.31 | 0.98 | 0.66 | 0.36 | 1.19 |
| 35-49 | 2.13** | 1.42 | 3.19 | 2.20** | 1.46 | 3.33 |
| Parity (reference = multiparous) |  |  |  |  |  |  |
| Primiparous | 1.46** | 1.10 | 1.94 | 1.41** | 1.05 | 1.89 |
| Women's education (reference = Primary or less) |  |  |  |  |  |  |
| Secondary |  |  |  | 0.94 | 0.61 | 1.47 |
| Tertiary |  |  |  | 1.23 | 0.73 | 2.08 |

|  | Restricted model |  | Complete model |  |
| --- | --- | --- | --- | --- |
|  | OR | 95% CI | OR | 95% CI |
| Household wealth quintile (reference = Poorest) |  |  |  |  |
| Poor |  |  | 1.01 | 0.61 1.69 |
| Middle |  |  | 0.91 | 0.53 1.55 |
| Rich |  |  | 0.85 | 0.48 1.48 |
| Richest |  |  | 1.32 | 0.71 2.47 |
| Ethnicity (reference = Kinh/Hoa) |  |  |  |  |
| Ethnic minorities |  |  | 0.61** | 0.40 0.93 |
| Region (reference = North Central) |  |  |  |  |
| Mekong River Delta |  |  | 0.82 | 0.54 1.25 |
| Red River Delta |  |  | 0.61** | 0.40 0.94 |
| Northern Midlands |  |  | 1.10 | 0.73 1.64 |
| Central Highlands |  |  | 0.54** | 0.35 0.84 |
| Southeast |  |  | 0.75 | 0.49 1.15 |

a\*\*: $p \leq 0.05$ , \*: $p \leq 0.10$
(Source: 2013-14 MICS)

To better understand the contrasts between urban and rural dynamics as well as correlates of CS, we will separately study women from each place of residence. The results are displayed in Table 3. They show two models: one concerning urban areas, and the other one concerning rural areas. In both models, maternal age at delivery over 35 is a major positive correlate of CS. Beyond this common phenomenon, distinct lines of socioeconomic and demographic cleavage operate in urban versus rural areas.

**Table 3.**
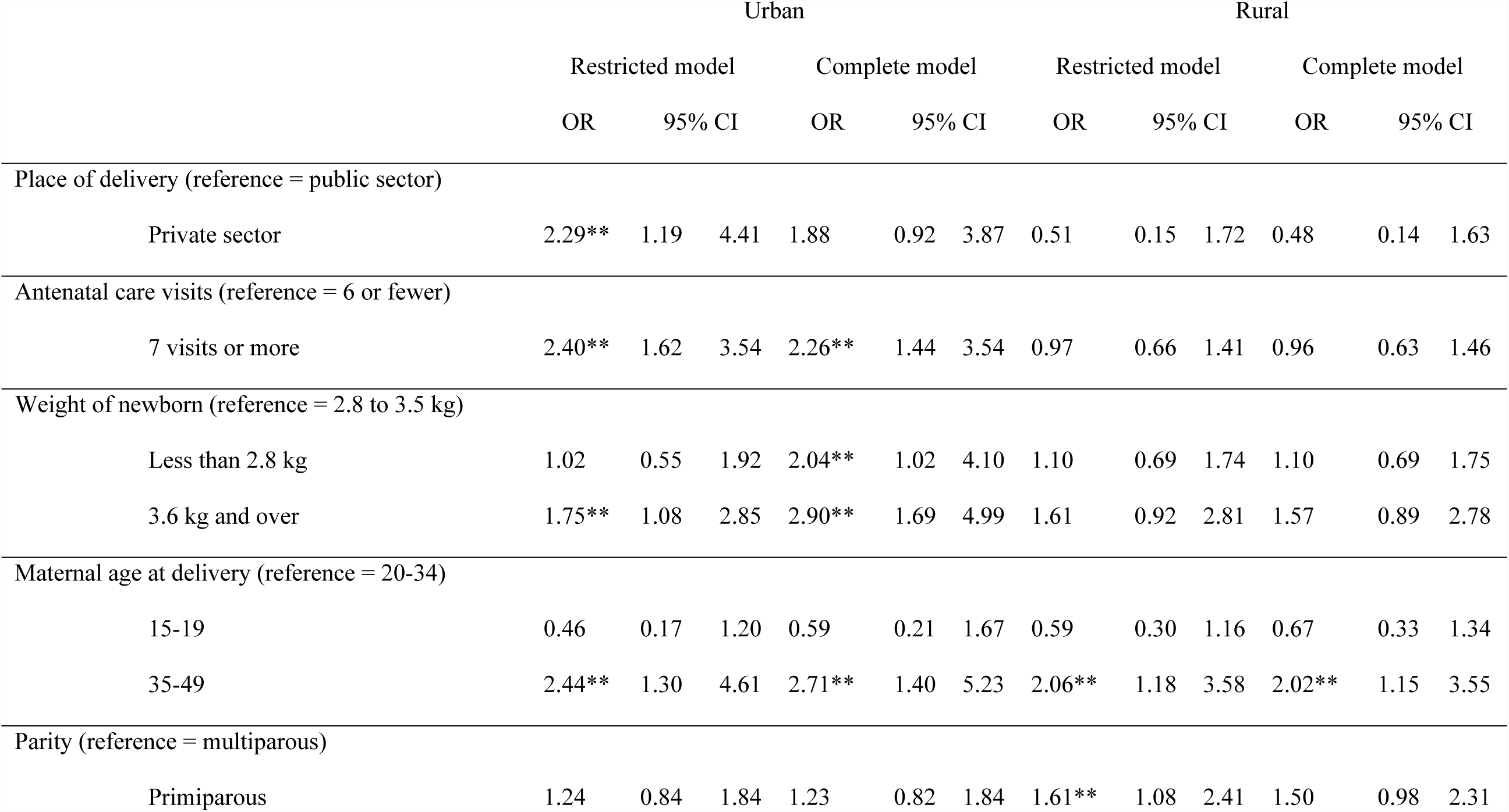
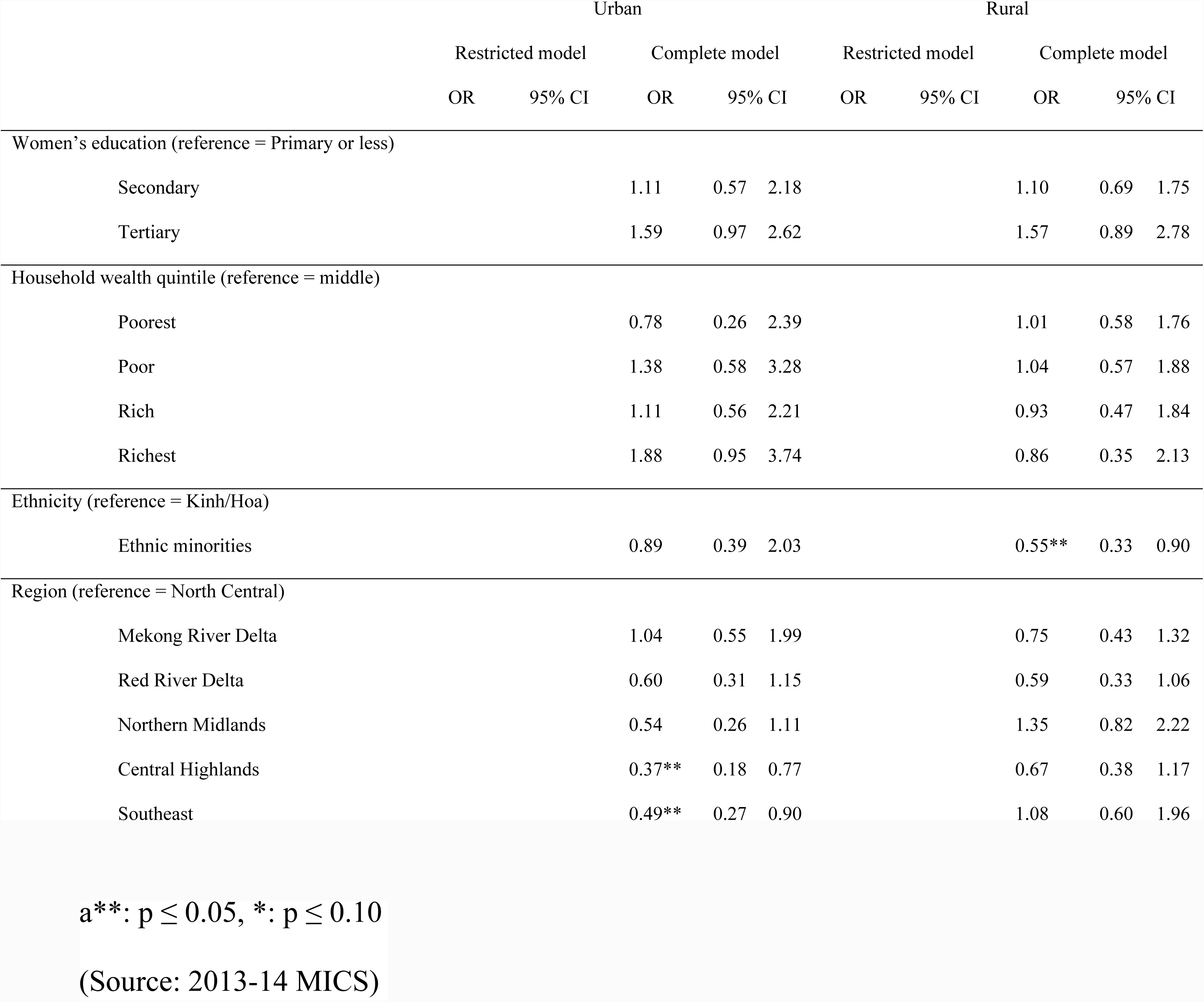
Odds of undergoing caesarean section (CS) for all women according to their place of residence: urban (n = 415) and rural (n = 935) areas^a^.

In urban areas, women are more than twice more likely to undergo CS when they have at least 7 antenatal care visits. They are also more than twice more likely to have CS when they deliver in the private health sector. Perception of one’s baby’s weight as above average also has a strong effect. When socioeconomic characteristics are taken into account, the distinction between private and public health sector disappears, whereas living in the Central Highlands or in the Southeast region appears linked to lower levels of CS. This suggests that the influence of private or public health sector is partly explained by contrasts between regions.

In rural areas, parity has a significant effect in the restricted model. Primiparous women are twice more likely to undergo caesarean section than multiparous women. This effect does not remain when sociocultural factors are taken into account. In the complete model, the odds of undergoing CS are almost halved for women belonging to minority ethnic groups. This suggests that the higher level of CS among primiparous women may be partly explained by the fact that they belong to the Kinh ethnic group, where fertility reaches lower levels.

## Discussion

The findings of this study confirm our primary assumption that the place of residence has a significant effect on CS practices in Vietnam in 2013-14. This outcome contrasts with the narrowing rural–urban gap in childbirth medicalization (13)(12). It updates the previous results showing a nonsignificant influence of urbanization on CS in the early 2000s (9). At the same time, despite growing levels of urbanization, nearly half of all CSs still occur in rural areas, as has been the case for the last two decades (33)(32)(12). This trend can be explained by the combination of doubling urbanization rates since 1997 and a more rapid increase of CS rates in rural areas than in urban areas.

The main determinant of CS is maternal age at delivery. This has also been reported in previous research in Vietnam (38)(36) and other countries (7)(8)(17)(10). On average, the mean age at childbearing is 24.7 years in Vietnam, and this indicator has remained stable between 24 and 24.8 years over the last decade (35). Our study reveals that a maternal age of over 35 years at childbirth more than doubles the likelihood of undergoing a CS and that this effect is stronger in urban areas, where childbearing is experienced slightly later than it is in rural areas.

To understand the factors leading to a CS, we have to take into account the circumstances of childbirth as well as the whole process of pregnancy. This need is underlined by the positive influence of a high number of antenatal care visits, which prevails in urban areas. This phenomenon, which is linked to high levels of antenatal ultrasound, may also be the consequence of pregnancy complications. It has been observed in Vietnam mostly in relation to prenatal sex selection and fear of birth defects (39). This trend has also been witnessed in eastern China, where it has been proven to be linked to a high level of CS practice (40).

A contrast exists in the factors associated with caesarean delivery in rural and urban areas. Various underlying social, demographic and economic rationales are involved. The perception of the weight of the newborn over or below average significantly increases the likelihood of undergoing a CS in urban areas, whereas it has no significant effect in rural areas. This result complements previous findings regarding periurban settings in Northern Vietnam, which showed no effect of the weight of the newborn on the mode of delivery (38). This greater use of CS in cases of macrosomia or low-weight newborns may be linked to the availability of services.

In addition, the influence of social networks in urban areas could be stronger than that in rural areas due to a higher level of instruction, higher level of exposure to the media and greater involvement of women in formal professional activities. Interestingly, women’s education level has no significant effect. The media hold power over healthcare facilities through the diffusion of information on their practices and results. Part of this power is used through social networks, by which public opinion is shaped. Women’s abilities to argue their cases and seek legal recourse in case of medical complications may act as a more powerful form of pressure on health staff in urban areas (41)(3). (25). The higher levels of human and social capital of women could make it more difficult for health personnel to resist women’s requests to undergo CS.

The highest rates of CS are observed among the richest household quintiles. This confirms a widespread trend in many countries (6)(3)(42). It also illustrates the persistence of inequalities in Vietnam despite some progress (43)(30). However, the household level of wealth has no effect after controlling for other sociocultural factors. This absence may be partly explained by social insurance coverage, which, despite lower levels of coverage in rural areas than in urban areas, covers 70% of the population (44). A positive link between health insurance and CS practice has been observed in neighboring China (45) and may apply to Vietnam despite problems with low protection levels (46), especially in rural areas (47).

In contrast with previous research in other low- and middle-income countries, our results show no influence of the private health sector after other sociocultural factors are taken into account (41)(8)(48). In Vietnam, the development of private healthcare facilities has undergone a major evolution following the Doi Moi reforms launched in the mid-1980s. However, the proportion of women delivering in this sector remains low. The role of the private health sector may be underestimated due to the offering of private services in public health facilities. A more in-depth investigation distinguishing between private and public services within the public health sector could provide more insights. Further explorations of our data show that the proportion of women who deliver in the private health sector varies widely across regions. As an example, the proportion is close to zero in the Northern Midlands and Mountain area but reaches 20% in the urban Mekong River delta.

Hence, urban areas appear heterogeneous across regions, and the pattern of this heterogeneity is unexpected. The lowest levels of CS are reached in the Central Highlands, which is understandable given the low level of equipment and the population density in this area. Surprisingly, a low level was reached in the highly urbanized and densely populated Southeast region, where indicators of medical equipment were much more favorable for CS. Further investigations show that delivery in the private health system and a high number of antenatal care visits are prominent factors of CS rates in this region, suggesting a complex combination of determinants.

This complexity is also illustrated by the fact that heterogeneity in CS rates depends on the region in urban areas but not in rural areas, where the key factor is ethnicity, which in turn is not relevant in urban areas. Women from minority ethnic groups are less likely to perform a CS regardless of the other characteristics taken into account in our study. This gap is widened by a lower level of birth in health facilities as well as a lower level of assistance by skilled attendants during delivery among women belonging to minority ethnic groups (34). It argues in favor of a sociocultural dimension of attitudes and opinions towards childbirth, which may involve interpersonal communication and transmission. Through our stepwise methodology, ethnicity appears to be a hidden factor.

CS determinants may combine with each other. Trends towards lower fertility in urban areas are in favor of higher levels of antenatal care attendance and CS use (35). Experience in other countries shows that in a context of reduced fertility, couples tend to be more willing to invest in the monitoring of pregnancy and caesarean delivery (17). This phenomenon should be distinguished from the concept of “precious pregnancies” attached to low-fertility couples, which has been subjected to criticism (49). A previous study performed in Vietnam showed that discussions with relatives also play a moderating role in helping women avoid CS (9). Such discussions may be more frequent in rural areas than in urban areas, where the family size is smaller (50). One heuristic concept capable of integrating the factors of CS may be “urban liveability”, which encompasses not only the physical setting but also social interactions and has been studied in relation to the social determinants of health in northern countries (51). Another factor worth exploring is the influence of the household registration system. In China, this system has proven to be more strongly linked to unmet long-term care needs than the place of residence (52). The question of whether similar effects on prenatal healthcare apply in Vietnam can be explored because the health sector is spatially divided for heath infrastructures and health insurance schemes.

The different CS rates between rural and urban areas may also be explained by different levels of healthcare equipment. In Vietnam, where the health system is pyramidal with a special status for main cities (53)(54), the two metropolitan areas of Hanoi in the Red River delta and Ho Chi Minh City in the Southeast region play key roles. The fact that more than half of the urban population lives in either the Southeast region (29.9%) or the Red River delta (23.1%) reveals the demographic weight of the two main metropoles (Hanoi in the Red River delta and Ho Chi Minh City in the Southeast region). At the other extreme, almost half of women living in rural areas reside in the Red River delta (25.5%) or the North Central region (22.0%).

The two main metropolitan areas in the country benefit from a concentration of highly equipped healthcare facilities in densely populated zones served by viable transport and road networks (28). This situation leads to a high number of deliveries within specialized healthcare services, as exemplified by the National Hospital of Gynaecology and Obstetrics in Hanoi, where more than 20,000 deliveries take place annually, with a CS rate of 48%^1^. The rural–urban divide is further strengthened by competition between health infrastructures following the “autonomization” policy launched in the 2000s, which spurs hospitals to make profits from investments (55). In urban areas where health personnel are more heavily subject to time pressure and overcrowded services, CS enables more predictable staff management and shortens the delivery duration (56). Public hospitals at the tertiary level are closely monitored (53)(54). These hospitals where CSs are performed (Dinh et al., 2012) play a pioneering role in the elaboration and implementation of health policies at the national level (57).

This study has limitations. First, we do not know the reason why the CS deliveries under study have been performed. In particular, we cannot identify medically indicated CS deliveries and those performed upon the patient’s request. Therefore, we can only uncover general trends. Second, we do not distinguish between several levels or types of urbanization; this type of analysis would require a large sample size. Third, the place of residence may not coincide with the place of delivery. Therefore, we capture the impact of the long-term influences of the context rather than the impact of possible adaptation through migration. Fourth, our statistical analysis provides indications of correlations rather than causal links. However, we are convinced that this study provides useful insights into the influence of urbanization on CS through highlighting its major determinants and suggesting a way to approach this complex phenomenon using existing data representative of the national level.

## Conclusion

The overall CS rate among the women who delivered in healthcare facilities in Vietnam is particularly high (29.2%) with regards to WHO standards (14). After controlling for significant characteristics, living in urban areas more than doubles the likelihood of undergoing a CS (OR = 2.31; 95% CI 1.79 to 2.98). Maternal age at delivery over 35 is a major positive correlate of CS. Beyond this common phenomenon, our study has shown contrasting models regarding the determinants of recourse to high levels of CS rates between rural and urban areas. This contrast suggests that actions to reduce unnecessary caesarean deliveries should be adapted to each context. Indeed, our results show the importance of taking into account not only medical and sociodemographic factors but also sociocultural determinants when designing programs to improve women’s childbirth conditions. It is the case of ethnicity, which needs to be addressed. This approach involves policies at many different levels regarding not only the regulation of the health sector and training of healthcare providers but also the sensitization of the entire population, with means appropriate to their conditions of living. Further research must be conducted to design such programs and to provide guidance on this complex issue.

## Acknowledgements

The authors acknowledge the Vietnam General Statistics Office (GSO) and Vietnam UNICEF for providing the underlying data that made this research possible, with special thanks to Ms. Nguyen Quynh Trang (UNICEF Vietnam) and Mr. Nguyen Dinh Chung (GSO). They also thank IRD and CEPED for their support.

## Footnotes

1 For more information, see Nguyen, T. H. P. (2016). *Nghiên cứu tÍnh hÍnh mổ lấy thai tại bệnh viện ph? sản trung ương từ tháng 3/2016 đến 5/2016 [Research on the situation of caesarean section in Central Hospital of Gynecology and Obstetrics from March 2016 to May 2016]* (Internship medical thesis). Ministry of Health, Central Hospital of Gynecology and Obstetrics, Hanoi.

